# *In vitro* activities of the tetrazole SHR8008 compared to itraconazole and fluconazole against *Candida* and *Cryptococcus* species

**DOI:** 10.1101/2020.01.15.908616

**Authors:** Lili Wang, Min Zhang, Jian Guo, Wenzheng Guo, Ni Zhong, Hui Shen, Wenjuan Wu

## Abstract

*Candida* and *Cryptococcus* are the main pathogens of clinical fungal infection associated with high morbidity and mortality. SHR8008 (in fact, this is the only official name in China and it is called VT-1161 by FDA) is a novel tetrazole agent that selectively inhibits fungal CYP51A compared to mammalian cytochrome P450 enzymes to achieve a better antifungal effect. The *in vitro* activities of SHR8008 and its comparators itraconazole and fluconazole were determined in 127 *Candida* and 50 *Cryptococcus* strains isolated from Chinese patients in the last 2 years by Invasive Fungal Infection Group. The MICs of SHR8008 and other triazoles were measured by the Clinical and Laboratory Standards Institute guidelines M27-E4. For *Candida* spp., SHR8008 (geometric mean MIC=0.078 μg/mL) was 6.5-fold and 11.2-fold more potent than itraconazole and fluconazole, respectively. There is a good correlation of MICs between SHR8008 and itraconazole/fluconazole. The MIC values of SHR8008 against *Candida glabrata* and *Candida tropicalis* were significantly lower than those of fluconazole, while for *Candida albicans* and *Candida parapsilosis*, the differences between SHR8008 and fluconazole were not statistically significant, either. For *Cryptococcus* spp., SHR8008 (geometric mean MIC=0.024 μg/mL) was 21.7-fold and 104.5-fold more potent than itraconazole and fluconazole, respectively. Against the seven *Cryptococcus neoformans* isolates with elevated fluconazole MICs (≥8μg/mL based on the MIC90 value for this azole), SHR8008 maintained potent activity, with MICs ranging between 0.031 and 0.5 μg/mL. The results showed that tetrazole SHR8008 was more promising in the treatment of *Candida* and *Cryptococcus* infection than itraconazole and fluconazole.

## Introduction

A growing number of immunocompromised patients have augmented morbidity of fungal infections, which range from easily treatable superficial type to life-threatening invasive infections(1, 2). The annual fungal infection incidences of common pathogens, including *Candida*, *Cryptococcus neoformans* and *Aspergillus*, have reached more than one in 10,000, and the incidence of *Candida* infections has risen to fourth place in nosocomial infections(3–5). According to the China Hospital Invasive Fungal Surveillance Net (CHIF-NET) study, *Candida albicans* was the most common species (44.9%), followed by the *Candida parapsilosis* complex (20.0%), *Candida tropicalis* (17.2%), and the *Candida glabrata* complex (10.8%), with other species comprising 3% of *Candida* isolates(6).

There are several approved clinical antifungal agents that have had some success in reducing the high mortality of invasive fungal diseases such as candidiasis and cryptococcosis(7, 8). The guidelines recommend that the treatment for cryptococcosis includes the use of fluconazole as the primary treatment for mild to moderate pulmonary infection, or for consolidation and maintenance after induction therapy, including intravenous administration of amphotericin B and flucytosine for cryptococcal meningitis or complex pulmonary disease(9). However, current treatment of cryptococcosis remains limitations due to the reduced fluconazole susceptibility, the side effects of amphotericin B and the availability of treatment in resource-limited situations, such as the lack of access to 5-flucytosine and the high cost of liposome amphotericin B(10–12). With the extensive use of azole antifungal drugs in clinical practice, the world’s shift in favor of non-albicans *Candida* species is troubling. *Candida glabrata* and *Candida tropical* exhibited generally high rates of resistance to fluconazole. The emergence of multidrug-resistant *Candida albicans* and *Candida auris* poses a threat to global health(13–15).

The safer, more specific and more effective antifungal agents are needed. SHR8008 (in fact, this is the only official name in China and it is called VT-1161 by FDA), is a novel investigational tetrazole, which targets the biosynthesis of ergosterol by selectively inhibiting fungal CYP51. SHR8008 has a lower affinity for heme-iron and a greater affinity for the fungal CYP51 polypeptide than current azole drugs. As a result, SHR8008 more potently inhibits fungal CYP51 than current azoles and less potently inhibits host cytochrome P450 enzymes, resulting in greater CYP selectivity. Therefore, SHR8008 avoids toxicity and drug interactions that occur with the cross-reaction of the azole with the human cytochrome P450 enzyme(16, 17). Moreover, SHR8008 has shown promising preclinical potential in the treatment of a wide range of fungal diseases, including dermatomycosis, trypanosomiasis, mycosis, coccidiomycosis and so on(18–22). Thus, the objective of this study was to evaluate the antifungal effect of SHR8008 systematically and to compare it with the triazoles itraconazole (ITC) and fluconazole (FLC) against common fluconazole-sensitive or resistant *Candida* and *Cryptococcus* strains isolated from Chinese patients in the last 2 years by Invasive Fungal Infection Group (IFIG).

## Results

### All *Candida* species

SHR8008 MICs ranged from 0.016-4 μg/mL against 127 *Candida* isolates, with MIC_50_ and MIC_90_ values of 0.031 and 1 μg/mL, respectively. As shown in Figure 1, SHR8008 MICs using the 50% inhibition endpoint were towards the lower end of the concentration ranges tested for the majority of isolates. The SHR8008 MICs were also lower than those of FLC for all *Candida* isolates (Table 1) and this activity was maintained against FLC-resistant predominantly isolates. Based on GM MIC, SHR8008 (GM MIC=0.078 μg/mL) was 6.5-fold and 11.2-fold more potent than ITC and FLC, respectively, and the differences between SHR8008 and FLC were statistically significant (Table 1 and Fig.1A). Good correlation between SHR8008 and ITC/FLC MICs (Pearson correlation coefficients of 0.7210 between SHR8008 and ITC, Pearson correlation coefficients of 0.6859 between SHR8008 and FLC; Pearson *P*<0.001; Figure. 2A and B) was observed, suggesting that any *Candida* strains having high MICs of SHR8008 also show high MICs of both ITC and FLC. SHR8008 showed similar good activity against *C. guilliermondii*, *C. krusei* and *C. lusitaniae*. Then, The SHR8008 MICs were somewhat higher against those of ITC or FLC-resistant *Candida* isolates, with MICs ranging between 0.016 and 4 μg/mL (Fig. 3). The MICs of SHR8008 were somewhat higher against the *C. glabrata* isolates (range, 0.125 to 4 μg/mL; MIC_50_ and MIC_90_, 0.5 and 2 μg/mL, respectively). In contrast, the other isolates were inhibited by lower concentrations of SHR8008, as reflected by lower MIC ranges and MIC_50_ and MIC_90_ values. MIC values of SHR8008 against *C. glabrata* and *C. tropicalis* were significantly lower than those of FLC (Fig. 1B and Fig. 1C), while for *C. albicans* and *C. parapsilosis*, the differences between SHR8008 and FLC were not statistically significant, either (Fig. 1D and Fig. 1E).

**Table 1.** SHR8008, itraconazole and fluconazole MICs against all *Candida* isolates (n=127)

| Species | Drug<sup>a</sup> | MIC Range (µg/mL) | GM MIC<sup>b</sup> (µg/mL) | MIC<sub>50</sub><sup>b</sup> (µg/mL) | MIC<sub>90</sub><sup>b</sup> (µg/mL) |
| --- | --- | --- | --- | --- | --- |
| Total (n=127<sup>c</sup>) | SHR8008 | 0.016-4 | 0.078 | 0.031 | 1 |
|  | ITC | 0.063-4 | 0.506 | 0.5 | 2 |
|  | FLC | 0.063->128 | 0.873 | 0.5 | 32 |
| *Candida glabrata* (n=32) | SHR8008 | 0.125-4 | 0.49 | 0.5 | 2 |
|  | ITC | 0.25-4 | 1.114 | 1 | 2 |
|  | FLC<sup>df</sup> | 1-64 | 3.589 | 2 | 64 |
| *Candida tropicalis* (n=30) | SHR8008 | 0.016-4 | 0.044 | 0.016 | 1 |
|  | ITC | 0.25-2 | 0.741 | 0.5 | 2 |
|  | FLC<sup>ef</sup> | 0.125->128 | 1.023 | 0.25 | >64.0 |
| *Candida albicans* (n=32) | SHR8008 | 0.016-0.5 | 0.037 | 0.031 | 0.125 |
|  | ITC | 0.125-2 | 0.317 | 0.25 | 1 |
|  | FLC<sup>e</sup> | 0.063-64 | 0.403 | 0.125 | 8 |
| *Candida parapsilosis* (n=29) | SHR8008 | 0.016-2 | 0.042 | 0.031 | 1 |
|  | ITC | 0.063-2 | 0.212 | 0.125 | 1 |
|  | FLC<sup>e</sup> | 0.125-32 | 0.333 | 0.25 | 16 |

<sup>a</sup> ITC, itraconazole; FLC, fluconazole
<sup>b</sup> GM, geometric mean MIC; MIC<sub>50</sub>, MICs at which 50% of the isolates are inhibited; MIC<sub>90</sub>, MICs at which 90% of the isolates are inhibited.
<sup>c</sup> In addition to those specifically listed, there are two *Candida guilliermondii*, one *Candida krusei* and one *Candida lusitanae* in 127 *Candida* isolates.
<sup>d</sup> CLSI clinical breakpoints: resistant $\geq 64$ (µg/mL); susceptible dose-dependent $\leq 32$ (µg/mL);
<sup>e</sup> CLSI clinical breakpoints: resistant $\geq 8$ (µg/mL); susceptible dose-dependent = 4 (µg/mL); susceptible $\leq 2$ (µg/mL).
<sup>f</sup> $P < 0.01$ versus SHR8008.

**Fig. 1.**
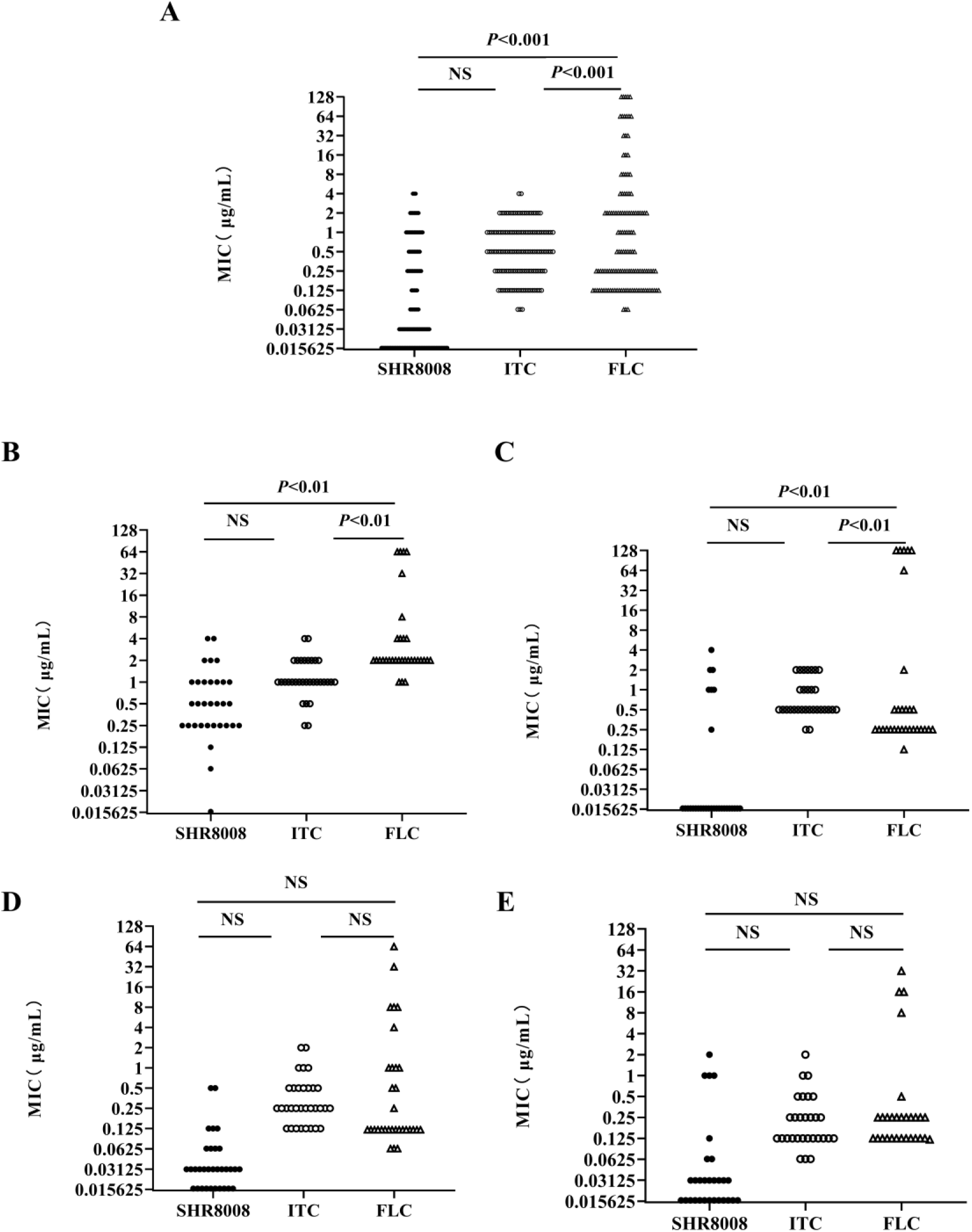
MICs of SHR8008, itraconazole and fluconazole for all *Candida* isolates tested. MICs of SHR8008, itraconazole and fluconazole determined by the CLSI method for all tested *Candida* isolates are plotted. (A) All *Candida* isolates. (B) *C. glabrata* isolates. (C) *C. tropicalis* isolates. (D) *C. albicans* isolates. (E) *C. parapslosis* isolates. *P*< 0.001, significant. NS, not significant; ITC, itraconazole; FLC, fluconazole.

**Fig. 2.**
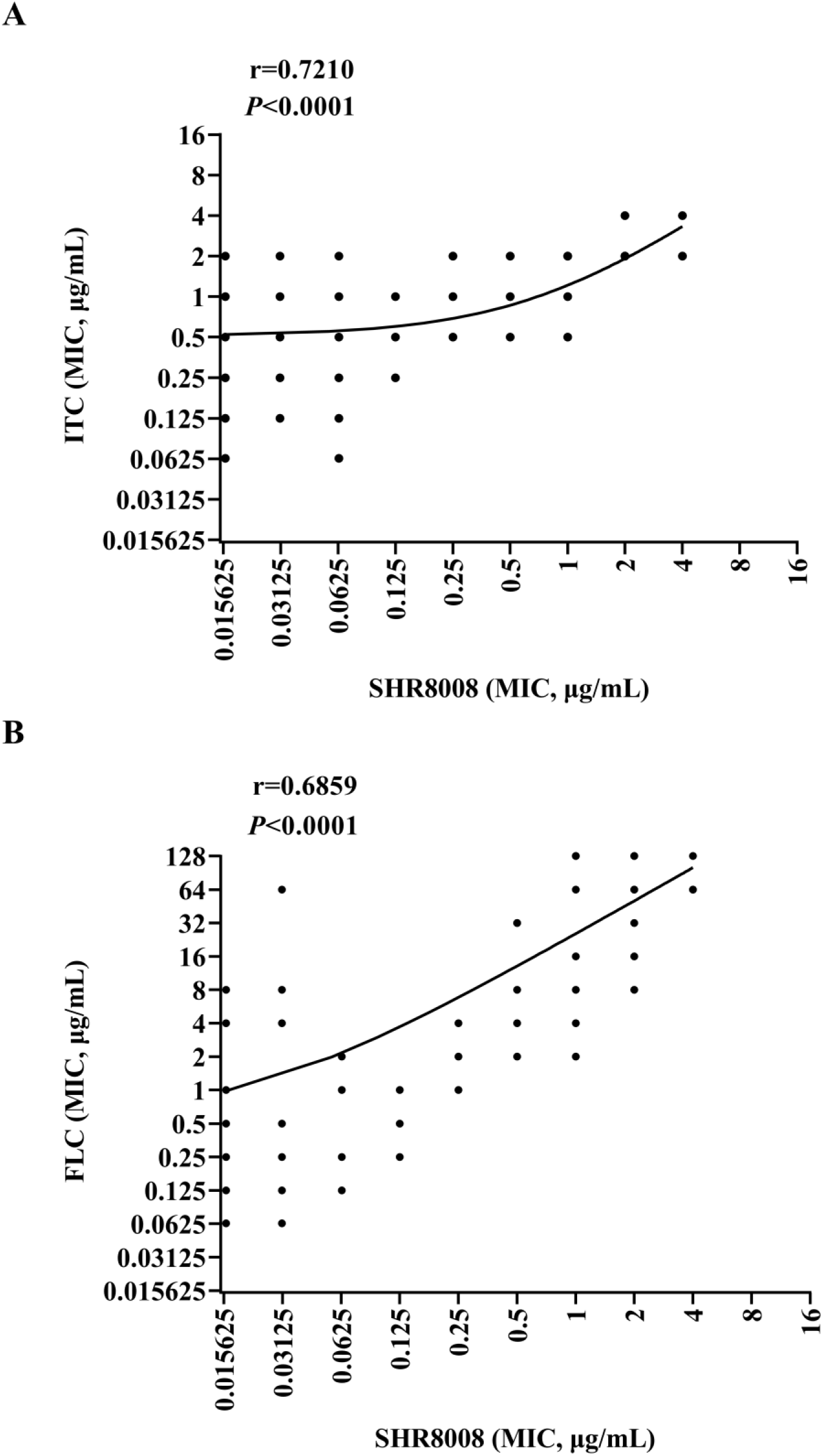
Individual MIC distribution and correlation. Relationship of MIC distribution for all *Candida* isolates between SHR8008 and itraconazole (A), fluconazole (B) is shown. ITC, itraconazole; FLC, fluconazole. R, Pearson correlation coefficient (| r |=1.0, completely correlated; | r |=0.8~1.0, highly correlated; | r |=0.5~0.8, significantly correlated. | r |=0.3-0.5, low correlation; | r |<0.3, weak correlation; R =0, no correlation); *P*<0.0001, the two are significantly (strongly) correlated.

**Fig. 3.**
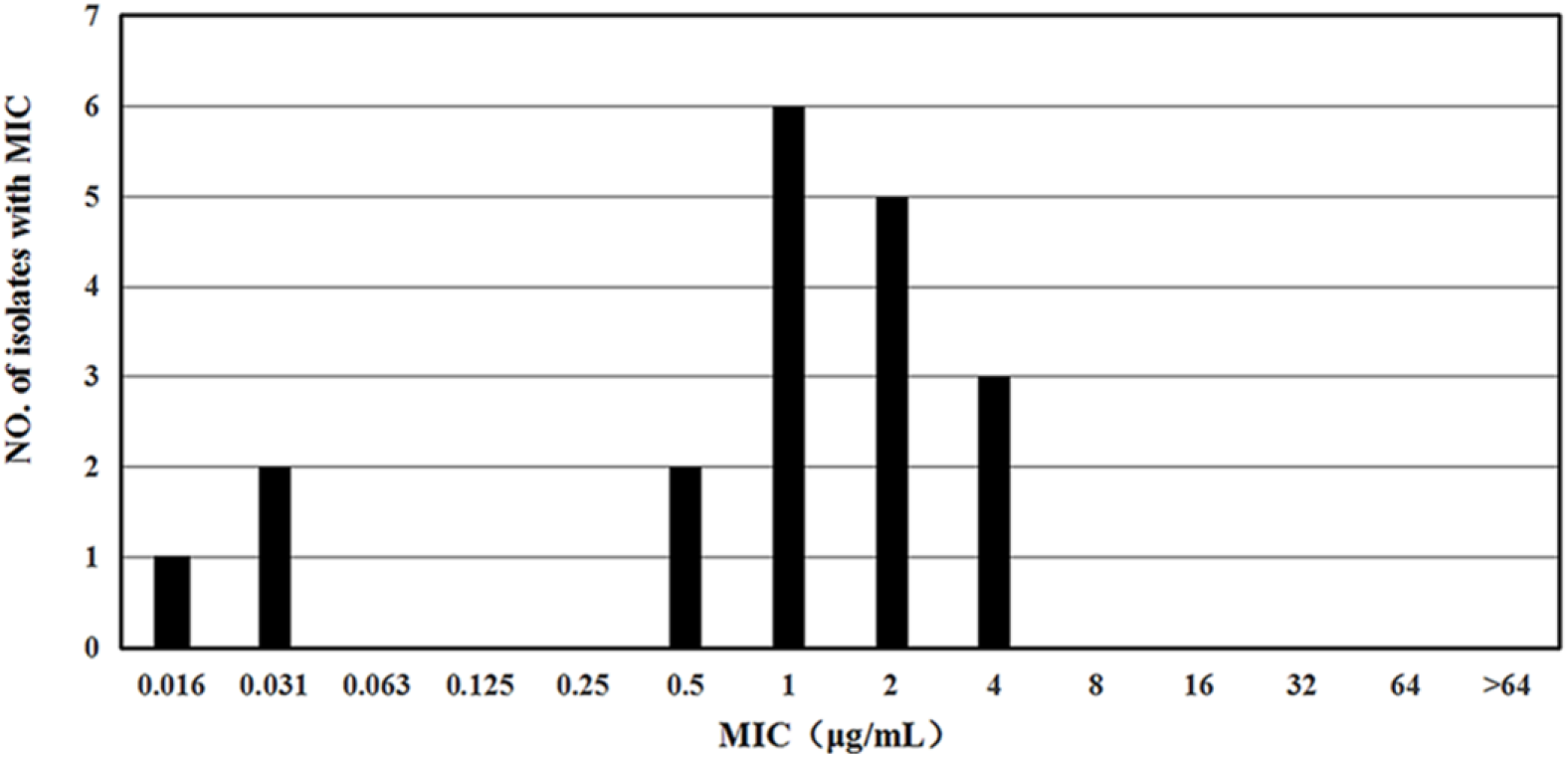
Distribution of SHR8008 MICs against all fluconazole-resistant *Candida* isolates (n=19).

### *Candida glabrata* and *Candida tropicalis*

SHR8008 demonstrated *in vitro* activity against 32 *Candida glabrata* isolates and 30 *Candida tropicalis* isolates tested in this study (Fig. 4A and Fig. 4B). All *C. glabrata* and *C. tropicalis* isolates were inhibited by SHR8008 at concentrations of ≤4 μg/mL after 24 h of incubation. The MIC_50_ of SHR8008, ITC and FLC against *C. glabrata* isolates were 0.5, 1, 2 μg/mL, respectively. The MIC_50_ against *C. tropicalis* isolates were 0.016, 0.5, 0.25 μg/mL, respectively. Based on GM MIC values read at 24 h, SHR8008 against *C. glabrata* was 2.3 fold and 7.3 fold more potent than ITC and FLC, respectively. And SHR8008 against *C. tropicalis* was 16.8 fold and 23.3 fold more potent than ITC and FLC, respectively. In *C. glabrata* and *C. tropicalis* isolates the MIC_90_ values of SHR8008 were 2, 1 μg/mL, respectively.

**Fig. 4.**
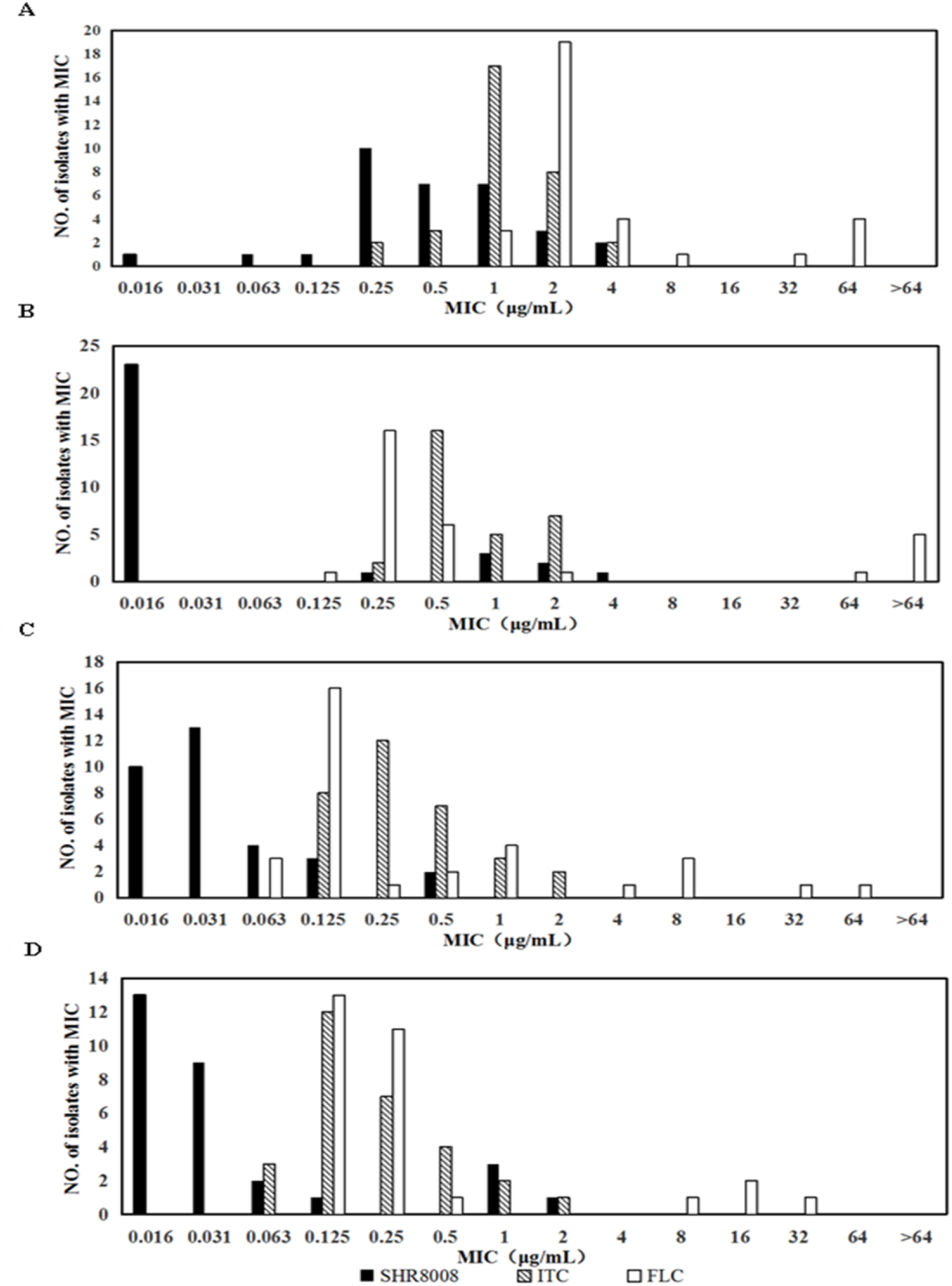
Distribution of SHR8008, itraconazole and fluconazole MICs against *Candida* isolates. (A) *C. glabrata* isolates. (B) *C. tropicalis* isolates. (C) *C. albicans* isolates. (D) *C. parapslosis* isolates.

### *Candida albicans* and *Candida parapsilosis*

SHR8008 demonstrated *in vitro* activity against 32 *Candida albicans* isolates and 29 *Candida parapsilosis* isolates tested in this study (Fig. 4C and Fig. 4D). All *C. albicans* and *C. parapsilosis* isolates were inhibited by SHR8008 at concentrations of ≤2 μg/mL after 24 h of incubation. The MIC_50_ of SHR8008, ITC and FLC against *C. albicans* isolates were 0.031, 0.25, 0.125 μg/mL, respectively. And the MIC_50_ against *C. parapsilosis* isolates were 0.031, 0.125, 0.25 μg/mL, respectively. The GM MICs of SHR8008 for *C. albicans* and *C. parapsilosis* were very low (0.037 μg/mL; 0,042 μg/mL) and MIC values of SHR8008 were ≤ 0.5 μg/mL for all *C. albicans* (Fig. 4C). For *C. albicans*, SHR8008 (GM MIC=0.037 μg/mL) was 8.6-fold and 10.9-fold more potent than ITC and FLC, respectively. For *C. parapsilosis*, SHR8008 (GM MIC=0.042 μg/mL) was 5-fold and 8-fold more potent than ITC and FLC, respectively. In *C. albicans* and *C. parapsilosis* isolates the MIC_90_ values of SHR8008 were 0.125, 1 μg/mL, respectively.

### *Cryptococcus* species

Overall, SHR8008 demonstrated potent activity against *Cryptococcus neoformans* with MIC values ranging between 0.016-0.5 μg/mL (Table 2). SHR8008 was more potent than ITC and FLC for each isolate, as evident by the lower MIC range and MIC_50_ and MIC_90_ values (Fig. 5). With the 50% inhibition endpoint, the SHR8008 MIC_50_, MIC_90_, and GM MIC values were 0.016, 0.125, 0.024 μg/mL, respectively. Based on GM MIC, SHR8008 (GM MIC=0.024 μg/mL) was 21.7-fold and 104.5-fold more potent than ITC and FLC, respectively, and the differences between SHR8008 and FLC were statistically significant. The MIC distributions for SHR8008, ITC and FLC against the *C. neoformans* isolates are shown in Fig.5. Against the seven isolates with elevated FLC MICs (≥8μg/mL based on the MIC_90_ value for this azole), SHR8008 maintained potent activity, with MICs ranging between 0.031 and 0.5 μg/mL. The SHR8008 MICs of tested isolates were directly plotted with MICs of ITC and FLC to visualize the relationship, indicating a certain correlation between them (Pearson correlation coefficients of 0.5443 between SHR8008 and ITC, Pearson correlation coefficients of 0.5139 between SHR8008 and FLC; Pearson P ≤0.001; Figure. 6A and B).

**Table 2.**
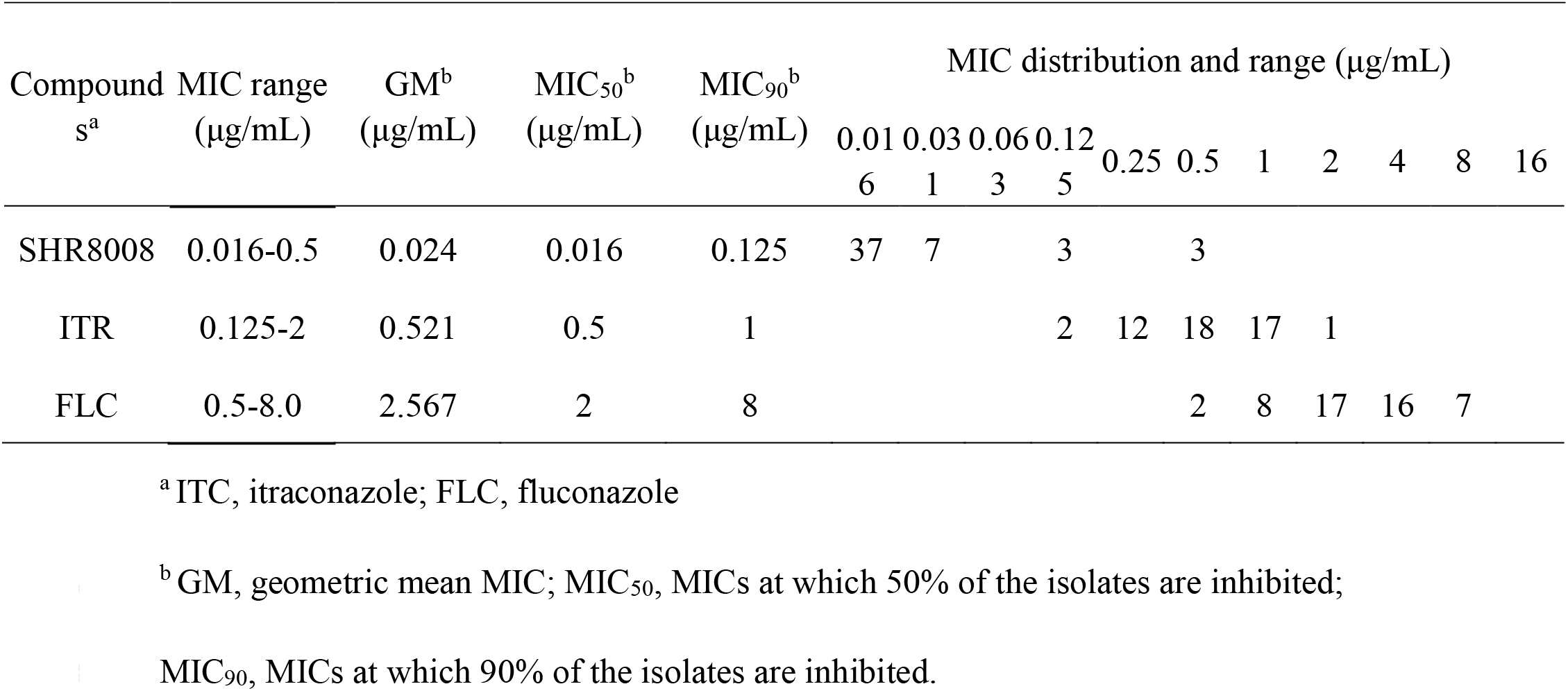
SHR8008, itraconazole and fluconazole MICs against *Cryptococcus neoformans* isolates (n=50)

| Compound<br>s<sup>a</sup> | MIC range<br>(µg/mL) | GM<sup>b</sup><br>(µg/mL) | MIC<sub>50</sub><sup>b</sup><br>(µg/mL) | MIC<sub>90</sub><sup>b</sup><br>(µg/mL) | MIC distribution and range (µg/mL) |  |  |  |  |  |  |  |  |  |  |
| --- | --- | --- | --- | --- | --- | --- | --- | --- | --- | --- | --- | --- | --- | --- | --- |
|  |  |  |  |  | 0.01<br>6 | 0.03<br>1 | 0.06<br>3 | 0.12<br>5 | 0.25 | 0.5 | 1 | 2 | 4 | 8 | 16 |
| SHR8008 | 0.016-0.5 | 0.024 | 0.016 | 0.125 | 37 | 7 |  | 3 |  | 3 |  |  |  |  |  |
| ITR | 0.125-2 | 0.521 | 0.5 | 1 |  |  |  | 2 | 12 | 18 | 17 | 1 |  |  |  |
| FLC | 0.5-8.0 | 2.567 | 2 | 8 |  |  |  |  |  | 2 | 8 | 17 | 16 | 7 |  |

<sup>b</sup> GM, geometric mean MIC; MIC<sub>50</sub>, MICs at which 50% of the isolates are inhibited;
MIC<sub>90</sub>, MICs at which 90% of the isolates are inhibited.

**Fig. 5.**
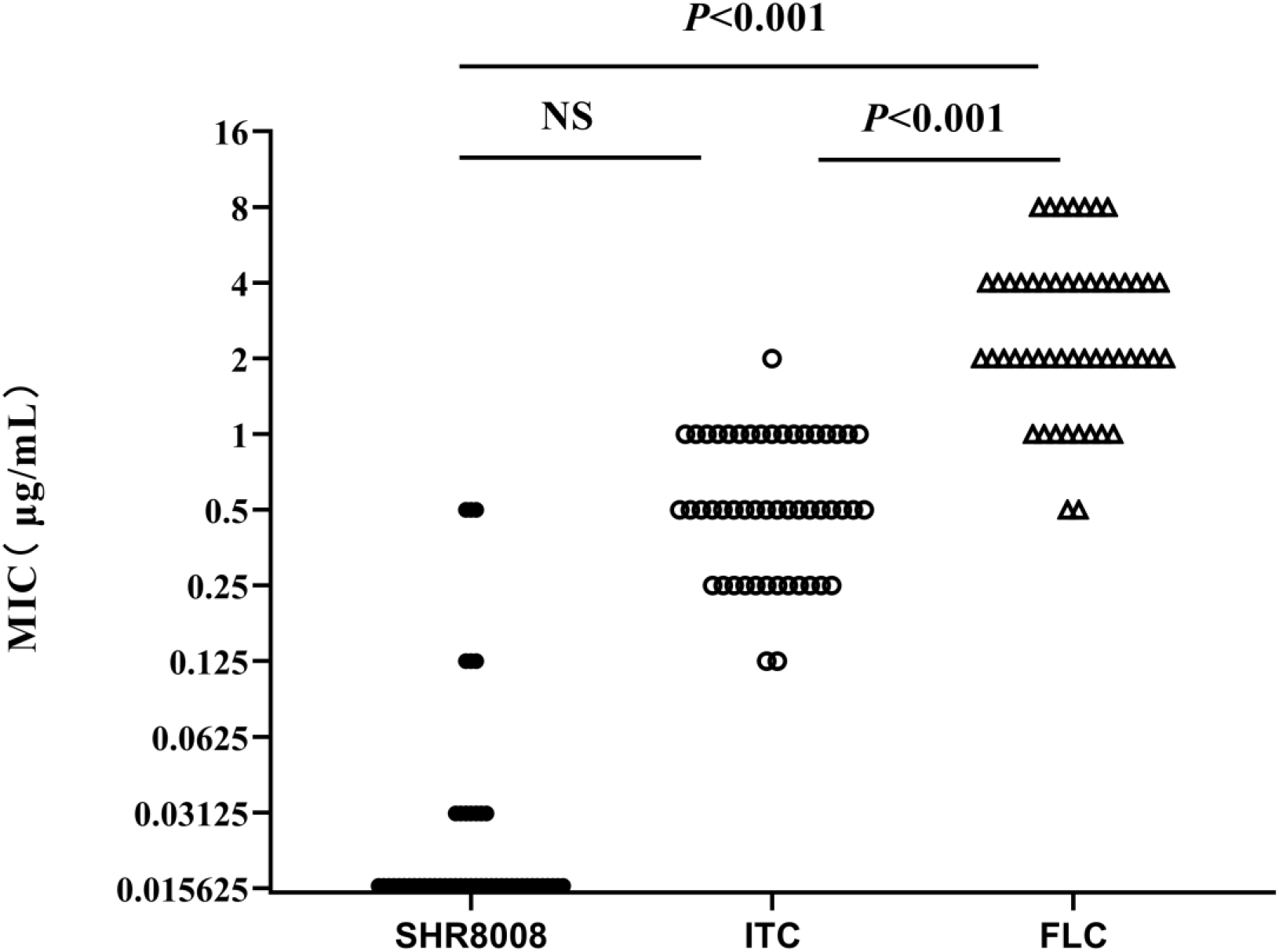
MICs of SHR8008, itraconazole and fluconazole for all *Cryptococcus neoformans* isolates tested. MICs of SHR8008, itraconazole and fluconazole determined by the CLSI method for all tested *Cryptococcus neoformans* isolates (n=50) are plotted. *P*< 0.001, significant. NS, not significant; ITC, itraconazole; FLC, fluconazole.

**Fig. 6.**
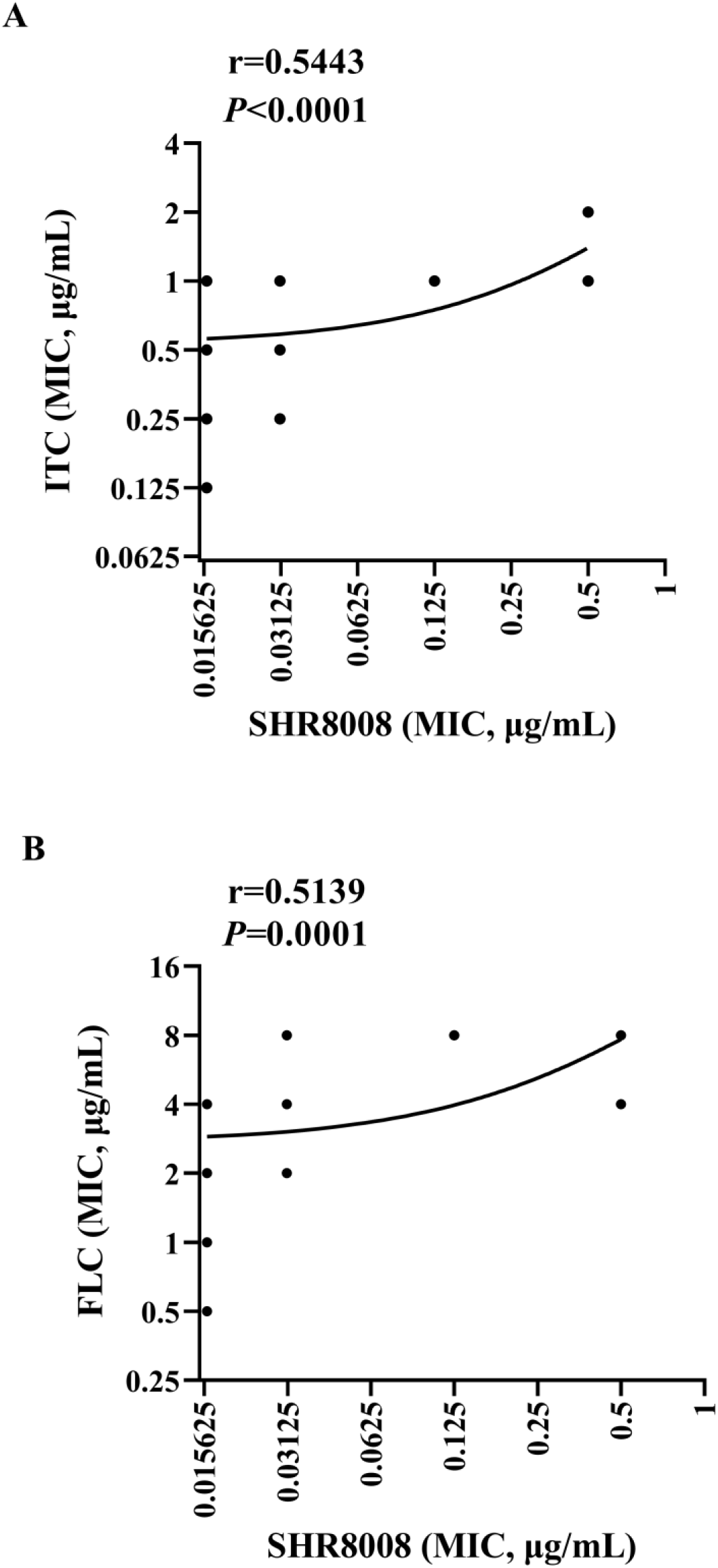
Individual MIC distribution and correlation. Relationship of MIC distribution for all tested *Cryptococcus neoformans* isolates between SHR8008 and itraconazole (A), fluconazole (B) is shown. ITC, itraconazole; FLC, fluconazole.

## Discussion

In this study, we systematically examined the *in vitro* susceptibility of yeast isolates obtained from IFIG to SHR8008, a novel tetrazole optimized for treatment, and SHR8008 was found to be a potent inhibitor of common fluconazole-sensitive or resistant yeast isolates. This tetrazole is more specific to fungal CYP51 than mammalian P450, so the potential for clinically relevant drug interactions is reduced compared to triazole agents that have been approved for use in humans(23).

The management strategies of recurrent vulvovaginal candidiasis (RVVC) is needed urgently, and the indication for SHR8008 in phase three clinical development is RVVC (https://www.mycovia.com). It is a debilitating, chronic infectious condition that affects millions of women. Primary symptoms include vaginal itching, burning, irritation and inflammation. RVVC is a global women’s health concern, which impacts the quality of life, to a degree comparable to asthma or COPD, and worse than diseases such as headache or migraine(24, 25). However, long-term maintenance suppressive fluconazole prophylaxis for RVVC frequently fails to cure the condition and serves only as an effective control measurement in many cases(26, 27). SHR8008 was shown to be effective and safe in the treatment of patients with recurrent vulvovaginal candidiasis(28). In our study, SHR8008 had more potential *in vitro* activity against *Candida* spp. than itraconazole and fluconazole and it was also promising in the treatment of invasive fungal infection. SHR8008 was found to be a tight-binding ligand and a potent inhibitor of CYP51 from the fungal pathogen *Candida* spp.(17).

We found that the MIC_50_ of SHR8008 against the *C. glabrata* isolates was 0.5 μg/mL, which was higher than those against *C. albicans*, *C. parapsilosis* and *C. tropicalis* isolates (MIC_50_ ≤ 0.031 μg/mL) (Table 1 and Fig. 4). Schell *et al.* described that the MIC_50_ values of SHR8008 were 0.12 μg/mL and 0.25 μg/mL for resistant *C. glabrata* and *C. krusei*, respectively(29). A previous study has reported that the *in vitro* and *in vivo* activity of SHR8008 was maintained against fluconazole-resistant *C. albicans* isolates(30). In the current study, this activity was also maintained against other *Candida* isolates with resistance to azole (Fig. 3). However, Nishimoto *et al.* found that SHR8008 may not be an ideal alternative in triazole-resistant *C. glabrata* infection, as it appears to be a substrate of the same efflux pumps associated with triazole resistance(31). Monk *et al.* found that azole resistance predicts reduced susceptibility to the tetrazole antifungal SHR8008, which is similar to this study(32). As shown in Fig. 2, a good correlation between MICs of ITC/FLC and SHR8008 was observed in a MIC distribution graph. MIC values of SHR8008 against *C. glabrata* and *C. tropicalis* were significantly lower than those of fluconazole (Table 1), so SHR8008 might make a breakthrough in the treatment of non-albicans infection.

*Cryptococcus* is a problematic opportunistic pathogen that causes invasive infection with high mortality. In particular, HIV-associated cryptococcal meningitis results in 150,000–200,000 deaths per year in sub-Saharan Africa(33–35). A clinical study showed that 30% of patients with cryptococcal meningitis receiving recommended therapy died by day 70(36). SHR8008 also demonstrated activity against *Cryptococcus* spp., with elevated itraconazole and fluconazole MICs (Fig. 5). MIC range of SHR8008 against *C. neoformans* was considerably low (0.016-0.5 μg/ml). In this study, we found a good correlation between SHR8008, itraconazole and fluconazole susceptibility for *Candida* spp., while a lower correlation was seen in *Cryptococcus* spp. (Fig. 6). The activity of SHR8008 against *Cryptococcus neoformans* was comparable to that previously reported the novel tetrazole VT-1129 (GM MIC=0.0271 μg/ml), although there was no direct comparison between them(37). These data suggest that SHR8008 may be a promising agent against *Cryptococcus* species. Further studies, for instance, the additional in vivo studies, including pharmacokinetic/pharmacodynamic evaluations and experimental models using isolates with different resistance profiles, are needed to determine how this investigational tetrazole may be used in the treatment of *Cryptococcus* infections.

In a murine model of Vaginal Candidiasis, Garvey *et al.* have demonstrated the efficacy of the clinical agent SHR8008 against fluconazole-sensitive and -resistant albicans, whereas fluconazole did not sustain efficacy 4 days post treatment(38). Recently, the first case of successful use of orally administered SHR8008 to treat RVVC has been published. SHR8008 exhibited higher levels of oral bioavailability and plasma protein binding than triazoles and showed excellent penetration into the vaginal and oral mucosa as determined in nonclinical models. In addition, there was also no evidence of an adverse effect of SHR8008 on liver function and SHR8008 further reduces the likelihood of problems arising from circulating drug substance(28). Thus, although the clinical breakpoints for the compound are not yet known, the MIC values reported here likely represent clinically relevant antifungal potencies.

SHR8008 is now under clinical development. The *in vitro* data of SHR8008 in our study indicates its potential to become a new generation of drug candidate for *Candida* and *Cryptococcus* infection prevention and treatment.

## Materials and methods

### Antifungal agents

SHR8008 (VT-1161, powders >99% pure) was synthesized by Mycovia Pharmaceuticals, Inc (DURHAM, NC, USA), whereas itraconazole (99% pure; Shanghai Aladdin Bio-Chem Technology Co., LTD, Shanghai, China), fluconazole (99% pure; National Institutes for Food and Drug Control, Beijing, China) were procured from commercial sources. Three drugs are supplied by Jiangsu Hengrui Medicine Co., Ltd. (Lianyungang, China).

### Strains

A collection of 177 clinical yeast strains isolated from IFIG, Shanghai East Hospital, were used in the study, including 32 *Candida albicans*, 32 *Candida glabrata*, 29 *Candida parapsilosis*, 30 *Candida tropicalis*, two *Candida guilliermondii*, *one Candida krusei*, one *Candida lusitaniae* and 50 *Cryptococcus neoformans*. *Candida parapsilosis* (ATCC 22019) and *Candida krusei* (ATCC 6258) were also used as quality control (QC) strains. They were inoculated onto CHROMagar *Candida* or Sabouraud dextrose agar culture medium following the National Clinical Laboratory Procedures (4th Edition). According to the color of the growing colony, the VITEK-2 Compact YST card (bioMérieux) or MALDI-TOF MS (bioMérieux or Bruker) was performed for preliminary identification. All the strains were identified by MALDI-TOF MS (Autobio) in the Central Laboratories (Shanghai East Hospital) again, and the strains with inconsistent results were identified by rDNA sequence analysis of the internal transcribed spacer (ITS) region (ITS1 primer: TCCGTAGGTGAACCTGCGG; ITS4 primer: TCCTCCGCTTATTGATATG).

### Antifungals and susceptibility testing

Antifungal susceptibility testing of yeasts was performed by broth microdilution in accordance with the Clinical and Laboratory Standards Institute (CLSI) M27-E4 reference standard(39). As stated in the protocol, Roswell Park Memorial Institute (RPMI) 1640 culture medium, with glutamine, without bicarbonate, and with phenol red as a pH indicator (Thermofisher), with 0.165 mol/L 3-(N-morpholino) propanesulfonic acid (MOPS; Sangong) was used to incubate strains across serially diluted concentrations of each azole. The 0.25–0.5*10^3^ cells/mL inoculum suspensions of yeasts were obtained from 24 h (*Candida* spp.) or 48h (*Cryptococcus* spp.) cultured Sabouraud dextrose agar plates at 35°C. Inoculum concentrations were verified by quantitative culture. Three antifungal agents were dissolved in dimethylsulfoxide (DMSO). The drugs were prepared at the following concentrations: 320 μg/mL for SHR8008 and ITC and 1280 μg/mL for FLC. The solutions were diluted in RPMI medium and the final drug concentrations ranging from 16 to 0.016 μg/mL, except for fluconazole, which ranged from 128 to 0.125μg/mL. The results were read separately at 24 h (*Candida* spp.) and 72 h (*Cryptococcus* spp.) after incubation. MIC endpoints were determined visually as the lowest concentration of compound that resulted in a decrease of growth by 50% relative to that of the growth control (azole endpoint: SHR8008, itraconazole, and fluconazole). MICs were determined in duplicate for clinical isolates. Categorical results were interpreted following the CLSI M60 document(40).

### Data analysis

Statistical analyses of all the data were performed using GraphPad PRISM 8 software program and the results are reported as MIC range, geometric mean MIC (GM MIC), MIC_50_ and MIC_90_. The MIC_50_ and MIC_90_ values reported for SHR8008, itraconazole and fluconazole, were defined as the minimum drug concentrations required to inhibit 50% and 90% of the clinical yeast isolates tested, respectively. Differences in MICs, calculated following log_2_ transformation of individual MIC values, were assessed for significance by ANOVA followed by Tukey’s multiple comparison test. A *P* value of <0.05 was considered statistically significant. The correlation between SHR8008 MICs and those of the other antifungals was also assessed by Pearson correlation using log_2_-transformed MIC values.

## Acknowledgements

We would like to thank Jiangsu Hengrui Medicine Co., Ltd. (Lianyungang, China) for providing SHR8008 (VT-1161), itraconazole and fluconazole drugs.

This study was funded by National Natural Science Foundation (grant number 81971990), Key Discipline Construction Project of Pudong Health Bureau of Shanghai (grant number PWZxk201709), Municipal Human Resources Development Program for Outstanding Leaders in Medical Disciplines in Shanghai (grant number 2017BR032).

